# Multimodal music perception engages motor prediction: a TMS study

**DOI:** 10.1101/358507

**Authors:** Chelsea L. Gordon, Marco Iacoboni, Ramesh Balasubramaniam

## Abstract

Cortico-spinal excitability (CSE) in humans measured with Transcranial Magnetic Stimulation (TMS) is generally increased by the perception of other people’s actions. This perception can be unimodal (visual or auditory) or multimodal (visual and auditory). The increase in TMS-measured CSE is typically prominent for muscles involved in the perceived action (muscle specificity). There are two main classes of accounts for this phenomenon. One suggests that the motor system mirrors the actions that the observer perceives (the resonance account). The other suggests that the motor system predicts the actions that the observer perceives (the predictive account). To test these accounts (which need not be mutually exclusive), subjects were presented with four versions of three-note piano sequences: sound only, sight only, audiovisual, and audiovisual with sound lagging behind while CSE was measured in two hand muscles. Muscle specificity did not interact with modality in the flexor digiti minimi (FDM), but was reliably higher for the first dorsal interosseous (FDI) while subjects perceived the audiovisual version of the three-note piano sequences with sound lagging behind. Since this version of the three-note piano sequences is the only one that overtly violates experience-based expectations, this finding supports predictive coding accounts of motor facilitation during action perception.

## Introduction

Motor regions of the brain are traditionally defined by their primary role in motor control (i.e., coding goals, planning, coordinating and executing actions) but motor areas additionally play a role in the perception of others’ actions (e.g., Hari et al., 1998; Buccino et al., 2001; Aziz-Zadeh et al., 2004; Fadiga et al., 2005). A common measurement used for detecting motor activation is transcranial magnetic stimulation (TMS) -induced motor-evoked potentials (MEPs), which reflect the level of corticospinal excitability (CSE) at the time of stimulation. MEPs have high temporal resolution, allowing for a precise measure of activity modulation. Increased CSE is found during visual perception of actions (Fadiga et al., 1995) as well as auditory perception of actions (Aziz-Zadeh et al., 2004). This increase is thought to reflect the recruitment of the mirror neuron system (Gallese et al., 1996), which is active both during action observation and action execution for similar actions, suggesting its involvement in the understanding of others’ actions (e.g., Fogassi et al., 2005; Iacoboni et al., 2005; Kaplan and Iacoboni, 2007; Gallese, 2008; Kilner, 2011). Populations of mirror neurons have been uncovered in premotor cortex that discharge upon observation and execution of the same action (Rizzolatti et al., 2001; Ferrari et al., 2003). The increased excitability in action perception is additionally time-dependent and effector-specific relative to the action being observed. Gangitano et al., (2001) recorded MEPs from the first dorsal interosseous (FDI) muscle during observation of a cyclic hand movement, and found that at the time when the FDI muscle of the observed hand was most contracted, MEPs recorded from the observer’s FDI muscle were highest. When the muscle was least contracted, MEPs were lowest. Thus the cortical motor areas of an observer are recruited for motor simulation of others’ actions in synchrony with those actions, and this is specific to the same muscle involved in the action.

Motor activity during action observation is also referred to as motor resonance, due to its time-dependent and effector-specific nature. The motor system of the observer “resonates” with that of the actor, allowing the observer to use their own body to understand, from within, the action being performed. One unanswered question is whether action observation is an active and predictive top-down process or a more automatic, bottom-up process. Early accounts, such as the direct-matching hypothesis (Iacoboni et al., 1999), suggest that motor resonance arises by directly perceiving an action via automatic activation in the observer of the cortical areas that represent the execution of that action. More recent theories propose that the mirror neuron system may function as a prediction mechanism during observation of others’ actions (Kilner et al., 2007). This differs significantly from the traditional assumptions of simulation (Gallese & Goldman, 1998), where the body of the observer “resonates” with the observed action but does not actively predict the future states of the movements. Active prediction proposes that top-down mechanisms influence the increase in CSE during observation, as a way to follow along with and actively predict another’s movements. It may be that mirror system activity actually reflects this predictive process, as the brain uses what it knows about the motor system of the observed actor to project the future state of the actor’s body. Because this type of prediction is very similar to that in motor control, the same neural systems (i.e., the motor system, mirror system) will underlie this process. Essentially, an observer can predict the motor commands of an observed actor given the expectations about their goal, and the implemented kinematics of that movement can be predicted using the observer’s own motor system.

Increased activity in motor areas has been observed in pianists when listening to piano pieces and observing piano playing (e.g., Haueisen and Knosche, 2001; Bangert and Altenmuller, 2003; D’Ausilio et al., 2006; Haslinger et al., 2005; Meister et al., 2004; Bangert et al., 2006), suggesting involvement of the mirror neuron system during music perception. Researchers have thus proposed that music is not passively heard, but actively perceived as the expressive motor acts that caused the music and are instantiated in the mirror neuron system (Molnar-Szakacs & Overy, 2006; Wallmark et al., 2018). Music observation is also a good candidate domain for exploring prediction during action observation, due to its sequential, and thus predictive, nature. Candidi et al. (2012) found that when expert pianists observed a fingering error (a note played with the incorrect finger), CSE recorded from the muscle corresponding to the finger playing the note increased significantly compared to the correct fingering of the keys. Non-musicians who were visually trained to detect the errors did not show this muscle-specific increase in corticospinal excitability during the fingering errors. The authors conclude that the experience of musically trained pianists provides their brains with *simulative error monitoring systems*. In other words, when participants were observing the fingering error, a prediction error occurred, leading to an increase in motor system activation. Furthermore, Stephan et al., (2018) showed in a recent study that auditory cues from a learned melody led to increased activation of the muscle that plays the *following* note of the melody, suggesting an anticipatory process occurring during melody processing in music perception.

Unexpected fingering errors cause one kind of prediction error that reflects an error of intent: the observer assumes that the player will play with one finger, and this prediction is violated when the player uses another. There are also purely sensory errors that can give rise to prediction errors, such as a multimodal stimulus with a misalignment between the auditory and visual components. Sensory errors that are not tied to specific effector movement error may not use the motor prediction system, as the prediction might differ from motor control prediction. On the other hand, existing research (Schubotz, 2007) has found that prediction of non-human-created sensory states (i.e., pitch prediction, object prediction) also relies on the motor regions of the brain. This suggests that we may see increased predictive activity in motor areas during a sensory prediction error as well. We aimed in our study to explore CSE modulation when there is no human movement error, but the sensory consequences of observed movements are temporally misaligned, resulting in a sensory prediction error. Specifically, the auditory correlate to a visual piano key press is delayed 200 ms. As the incoming visual signal of the motor act is perceived, a prediction of the corresponding auditory consequence is made. When the auditory signal is delayed so that onset is 200 ms into the video, the sensory prediction is violated. If the motor system is involved in sensory prediction, this stimulus should increase CSE due to error detection. Systems that use predictive coding mechanisms work optimally by using low resources when predictions are closer to the actual observed state, and increasing resource use during large discrepancies or errors (Mlynarski & Hermundstad, 2017). If effector-specific motor regions are recruited for generating sensory predictions, we can expect to observe an increase of CSE in the observed muscle during sensory discrepancies, but not an increase in other muscles (effector-specificity). For example, if a participant is observing a key played with the actor’s index finger, we should see increased excitability recorded from the participant’s index finger muscle, but not from their pinkie muscle. Furthermore, while detection of fingering errors increases corticospinal excitability in experienced musicians only, the prediction error in our study should give rise to facilitation in both musicians and nonmusicians, as no training is needed to understand the relationship between the observed action and the timing of its sensory consequences (c.f. Candidi et al., 2012).

In addition, while it is known that both visual and auditory action observation leads to increased CSE, the differential influences of each of these modalities on their own remains unclear. It is also unclear whether multimodal action perception will lead to additive activity in motor cortex summing over both visual and auditory contributions to motor regions. If motor involvement is primarily a bottom-up, automatic phenomenon, we should expect additive effects of multimodal presentation on CSE. The active inference framework, however, predicts that multimodal presentation will not lead to increased CSE over single-modality, as the information is redundant for predictive purposes. Therefore, an additional objective of the present work was to explore the modulation of CSE during auditory, visual, and multimodal music perception to explore the potentially different effects of modalities on motor system activity.

## Methods

### Subjects

Forty subjects were recruited for this study (15 males, mean age = 25 years). Due to excessive noise in the EMG signal or a participant having difficulty maintaining wakefulness, we excluded two subjects. All subjects were right-handed and had normal or corrected-to-normal vision and no hearing impairments. Subjects were screened for contraindications for TMS and previous medical problems that would be risk factors for TMS. The UC Merced Institutional Review Board approved the study, and written consent was obtained for all subjects. The experiment took about one hour, and subjects received two research credits that can be used for credit in some undergraduate courses.

### Stimuli

Eight different stimuli videos were used in our design. The videos were filmed using an iPhone 7 Plus camera and were edited using iMovie. These consisted of video recordings of a three-note piano sequence played using the right hand. Half of the stimuli were recordings of the thumb finger playing one note, followed by the index finger playing the next note twice. The finger movement that produces this action involves the first dorsal interrosseus (FDI) muscle. The other half had the same pattern but played by the ring finger followed by the fifth digit (pinky). This second half was played one scale higher than the keys used in the first half, so that the auditory cue for each could be distinguished. The finger movement that produces this action involves the flexor digiti minimi (FDM) muscle. For each of these two patterns, we created auditory-only (video blacked out), visual-only (audio silenced), multimodal, and multimodal time-lag versions (henceforth called incompatible), using iMovie. The time-lag version was created by starting the audio for the video 200 ms after the visual stimulus began. Each video was played 10 times in a randomized order, leading to a total 80 stimulations during the experiment. TMS stimulation was triggered by our presentation software, Paradigm, at the time in which the index or ring finger began its first down press, 1.5 seconds into the video.

### TMS and EMG recording

Corticospinal excitability was measured by the peak-to-peak amplitude of motor evoked potentials (MEPs) recorded using electromyography (EMG) on two muscles of the right hand. Two bipolar surface electrodes were placed on the belly of the participants’ right FDI muscle. Two additional electrodes were placed over the FDM muscle. A ground electrode was placed on a bone near the elbow of the subject. In order to obtain optimal EMG signal, we abraded and cleaned the skin under the electrodes, and secured the electrodes with medical tape. A bandpass filter (50 Hz–1000 Hz) was applied to the EMG signal, which was digitized at 1024 Hz for offline analysis. MEPs were elicited by applying single-pulse TMS to the region of the left motor cortex that induced MEPs in both FDI and FDM. If a location that induced MEPs in both muscles could not be determined, we used the FDI hotspot and thus did not record MEPs from the FDM muscle. Pulses were delivered using a Magstim Rapid2 TM with an attached 70 mm figure-of-eight coil positioned over the optimal scalp location with the handle pointing backwards at 45° from the midline. The motor hotspot localization procedure was as follows. Subjects were fitted with a swim cap that was covered by a grid of dots 1cm apart. Optimal scalp position was determined by moving the coil in 1cm intervals until the location eliciting the best MEPs in both muscles was identified. We were unable to find the shared hotspot position for eight subjects, and thus only have data from FDI for these subjects. The optimal location was marked on the swim cap worn by the participant. Resting motor threshold was determined as the percent of machine output that produced 3 out of 6 MEPs of at least 50 mV peak-to-peak amplitude. The stimulation intensity during the experiment was set to 120% of a participant’s resting motor threshold. The coil was held steady at the optimal position throughout the experiment. The interpulse interval between each stimulation was between 9 and 10 seconds. Subjects were instructed to keep their head still and remain relaxed with their right hand on their lap for the duration of the experiment, while attending to the videos as they appeared.

## Results

The EMG data was exported from Visor2 (ANT Neuro), and we ran a custom Python script to extract MEPs (peak-to-peak amplitudes). We also calculated area under the curve, but as these values correlated over 98% with the peak-to-peak amplitudes, we did not use both measures. In order to use inter-individual comparisons, Z scores were calculated for each participant. Trials in which MEP amplitudes were larger than 2.5 standard deviations from the mean and those less than 50 µV were excluded as outliers. Less than 5% of all data were excluded. Statistical analyses were carried out in R.

A repeated-measures analysis of variance (ANOVA) was conducted on MEPs from each muscle to assess the significance of the effect of our experimental conditions on the MEP amplitudes. We had a repeated-measures 2×4 design with two muscle congruence conditions (congruent was equal to 1 if the muscle that an MEP was recorded from was the same as the one observed in the trial, and equal to 0 if not) and 4 modality conditions (auditory, visual, multimodal, and multimodal incompatible).

In the ANOVA for the MEPs obtained from FDI, there was a significant main effect of modality (df = 3; F = 3.52; p = .01), wherein the incompatible stimuli induced significantly larger MEPs than the other conditions. We also observed an interaction between muscle congruency and modality (df = 1, 3; F = 4.93, p < .01), meaning that we did see muscle-specificity in MEP modulation in some conditions but none or less in others. We observed no additional main effect of muscle congruency. Post-hoc multiple comparisons using Tukey’s honest significant differences revealed that the incompatible congruent condition produced larger MEPs than the incompatible incongruent condition (t = .391, p < .05), while there were no significant differences between the congruency conditions in the other modalities.

In the ANOVA for the MEPs obtained from FDM, there was a significant main effect of muscle congruency (df = 1, F = 4.33, p < .05) indicating that we do see muscle-specificity in MEP modulation. Modality was marginally significant (df = 3, F = 2.23, p = .08), with the incompatible condition resulting in larger MEPs than the other conditions. We did not obtain an interaction between modality and muscle-specificity.

Average normalized MEP amplitudes for each modality condition and congruence condition can be seen in Figure 1. Overlaid example MEPs from the two congruence conditions in the incompatible modality are given in Figure 2.

**Figure 1.**
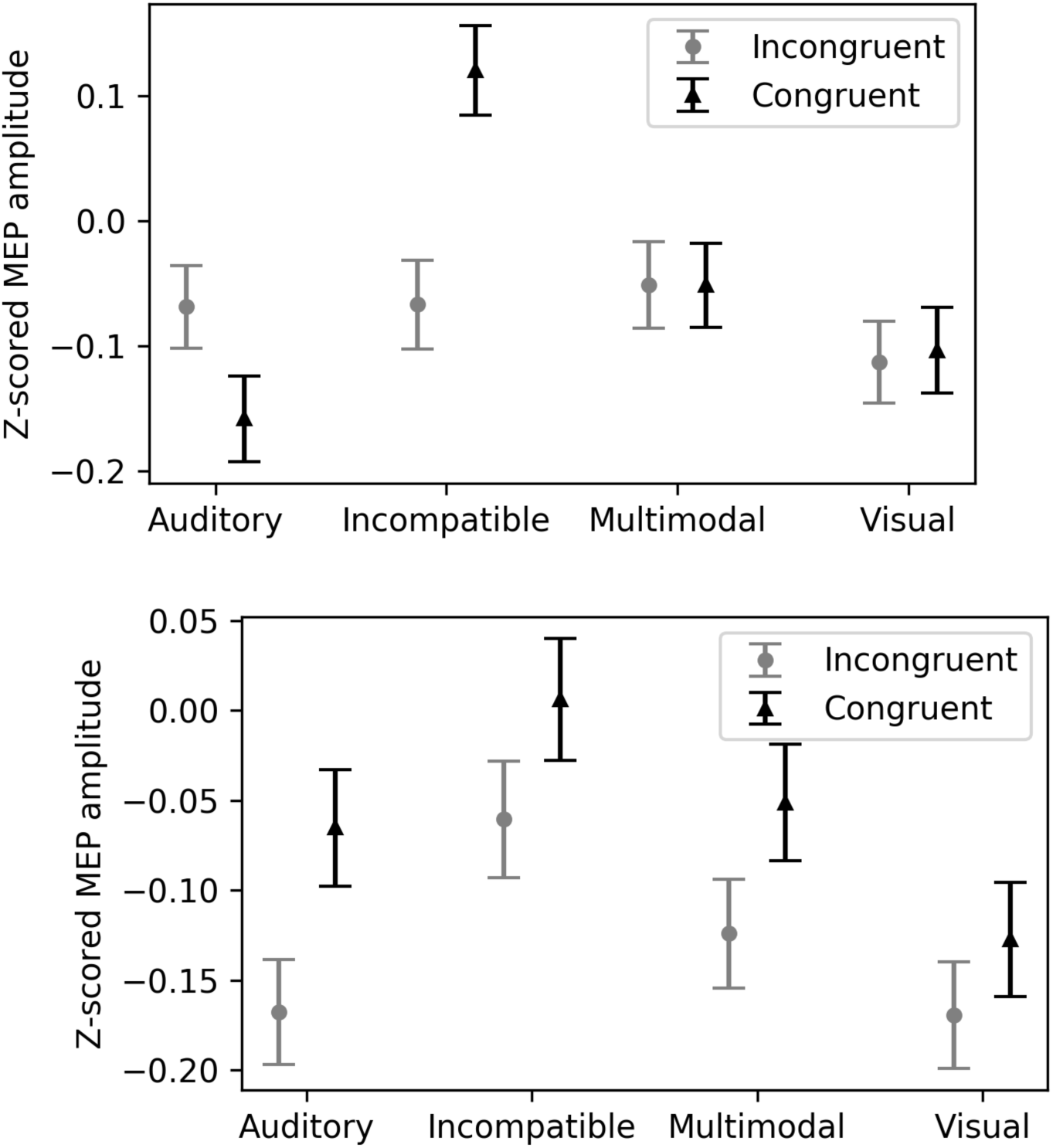
Z-scored MEP amplitudes for each condition and congruence condition. Data from all subjects. Vertical bars denote standard error of means. A) MEPs recorded from FDI. Motor evoked potentials in the incompatible incongruent condition show the largest facilitation. B) MEPs recorded from FDM.

**Figure 2.**
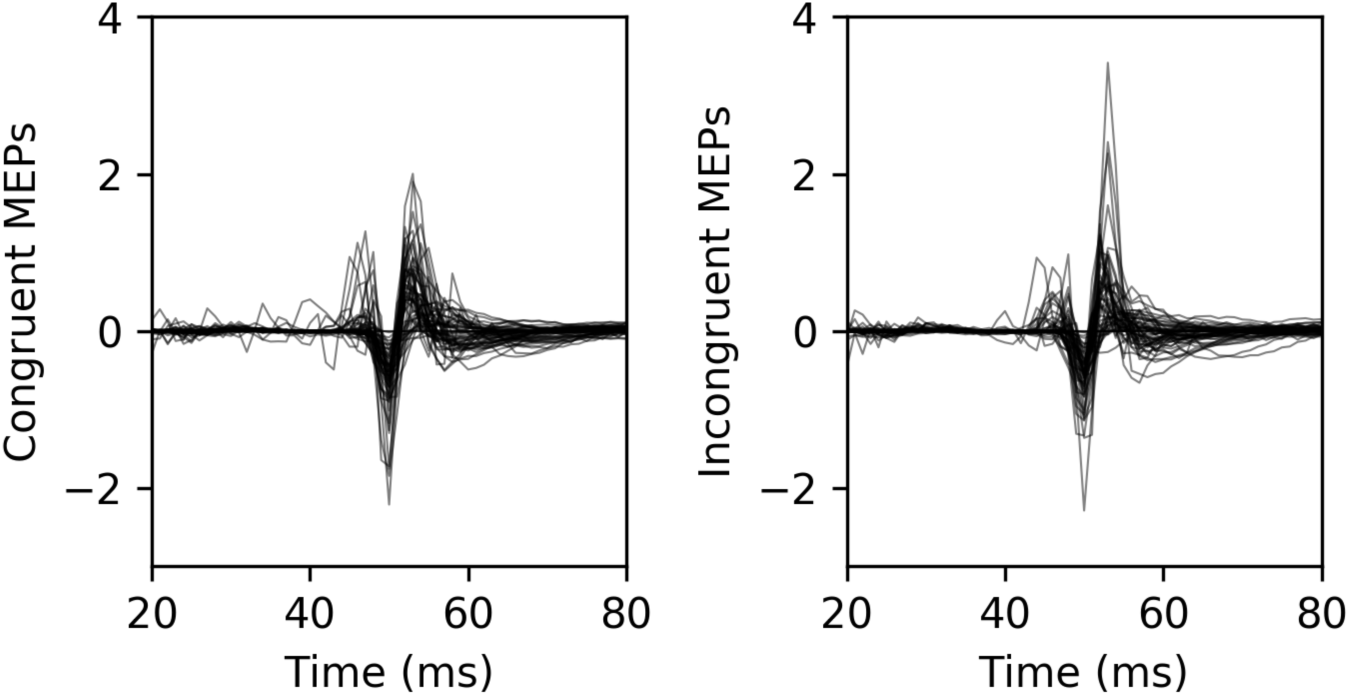
Representative example MEPs from 8 subjects during the incompatible trials. The left image contains MEPs from congruent incompatible recordings, and the right image contains MEPs from incongruent incompatible recordings.

To explore whether MEPs decreased as participants were exposed to more stimuli and whether this varied per modality, we ran a generalized linear model with subjects as a random effect and modality and trial number as fixed effects. MEPs decreased as trial number increased (df = 1, 1345, F = 7.87, p < .01), but we did not see a significant interaction between modality and trial number (F = .41, p ≈ .75). The linear trend for each modality by trial number is shown in Figure 3.

**Figure 3:**
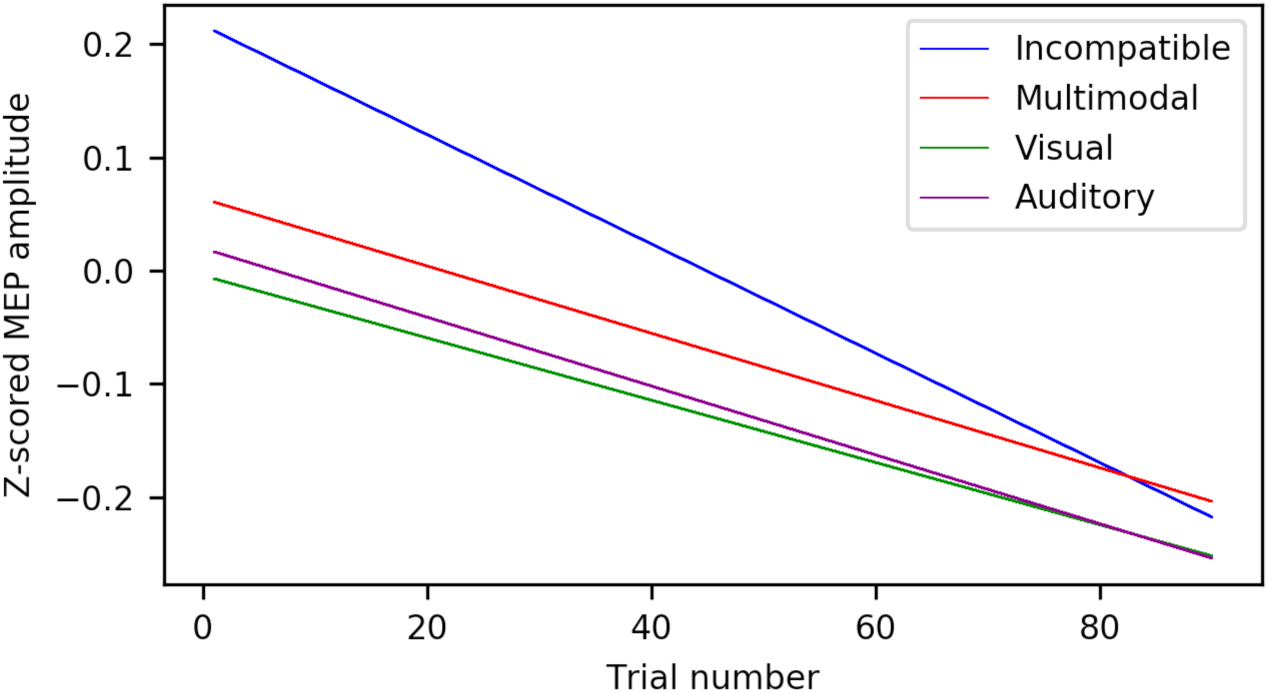
Linear fits for the modulation of MEP amplitude over number of trials completed.

## Discussion

In this study, we tested whether the increased in CSE during action perception is driven by bottom up resonance or by top down predictive coding. The two main tests we devised for these alternative hypotheses were as follows. Increased muscle specific CSE for the multimodal compatible stimuli, compared to the unimodal would support the bottom up resonance mechanism. A selective increase in muscle specific CSE for the multimodal incompatible stimuli, that violate sensory expectations, would support the top down, predictive coding hypothesis. The results support the latter, by showing an interaction between congruency and modality in the FDI, with larger MEPs for the incompatible congruent condition compared to the incompatible incongruent condition.

Corticospinal excitability increased the most for the incompatible incongruent stimulus, suggesting that the sensory error was detected and processed in motor regions. Prior work (Candidi et al., 2012) has shown that the motor system in musicians enables simulation of observed piano playing, and that activation increases when a fingering error by the pianist is observed. Here we provide evidence that motor activity also increases during the observation of a non-movement-related sensory error during action observation, which is present even in our participant pool, comprised primarily of non-musician subjects. The delayed onset of the auditory component resulted in a sensory prediction error, and a corresponding increase in corticospinal excitability. This provides evidence for a general predictive process taking place in motor areas of the brain at multiple levels, from intention prediction to sensory consequence prediction.

Active inference or “action-oriented predictive processing” has gained much interest over the last few years as a potential framework of how the brain instantiates perception, action, and cognition (Clark, 2015; Friston et al., 2011). Instead of considering the brain as a passive processor of bottom-up sensory information, these theories suggest that the brain is undergoing top-down active inference in order to predict incoming sensory information. Sensory information that is received provides feedback for top-down predictions to adjust predictive models in order to decrease prediction error in the future. Under this framework, prediction happens at multiple levels. At each level, generative models are created to predict information about the upcoming state of a lower level. Generative models calculate a prediction error based on a comparison of expected to actual sensory state. The prediction error is sent up the hierarchy, so that top-down mechanisms can adjust future predictions. This recurs until prediction error of the system is minimized. An important conceptual distinction in active inference theories is that motor processing is no different from sensory processing, as both are involved in top-down processing/prediction. A resulting idea from this is the existence of a single action-perception process that attempts to predict sensory input from all modalities. For action, the modality being predicted is proprioceptive input. The primary goal is to minimize surprise and thus minimize prediction errors.

In active prediction, neurons that are typically known to represent particular actions also represent the causes of sensory input (the same idea underlies ideomotor theory; Prinz, 2005; Hommel, 2013). In other words, perception and action share a common neural code. As such, Friston et al., (2011) suggest that the mirror neuron system can also be explained with active inference and predictive coding. Active inference implies a circular causality, whereby actions are deployed in order to fulfill predictions prescribed by perception, which updates these predictions using information obtained via actions. During action observation, the same process is instantiated, but without the corresponding proprioceptive feedback that occurs during action. This means that the same neuronal ensembles that encode an action during movement will encode that same action during observation. This naturally allows for the formation of mirror neurons, which will underlie this predictive process (Kilner et al., 2007) for both action observation and action execution. Neurons with mirroring properties have now been described in multiple systems of the primate brain. Beyond the original findings in fronto-parietal circuits for grasping (Rizzolatti & Craighero, 2004), mirroring response have been recorded in dorsal premotor and primary motor cortex for reaching movements (Dushanova & Donoghue 2010), in the lateral intraparieal area LIP for eye gaze (Shepherd et al. 2009), in the ventral intraparietal area VIP for touch (Ishida et al. 2010), and in human SMA and medial temporal cortex for grasping actions and facial expressions (Mukamel et al, 2010). This pervasive mirroring machinery seems ideal for generating predictive models during action observation. Higher-level generative models will make predictions about intentions and goals, while levels lower in the hierarchy will be involved in prediction of observed low level muscle movements.

This kind of predictive mirror neuron system can explain our results as well as the increased motor activation during fingering errors reported in Candidi et al. (2012), where error detection caused increased activation in predictive models to account for perceived error. When the visual component of the incompatible stimulus begins with no auditory counterpart, the prediction is that the given trial is a visual stimulus only trial. At the auditory component onset, this prediction is violated and there are misaligned sensory representations of an ongoing observed action. Furthermore, as participants receive repetitive exposure to all stimuli, corticospinal excitability decreases, suggesting a decreased prediction error that corresponds to learning. Though our model did not find a significant interaction between trial number and modality, visual examination of Figure 3 shows a slightly steeper downward slope for the incompatible condition relative to others. A longer experiment may be needed to detect whether there are differences in slopes for incompatible trials relative to other trial types, as in this experiment each stimulus was only seen 10 times. Future studies may also manipulate experimentally the number of trials that violate expectations within experimental blocks to further test the predictive coding hypothesis, as previously done for action preparation (Bestmann et al 2008).

In summary, sensory error detection during action observation leads to increased activation in the motor system. This facilitation likely results from prediction error caused by a mismatch between expected sensory consequence and actual sensory input. Sensory prediction errors may be generated in motor regions, and potentially rely on mirror neurons for this predictive process. This suggests that reconsidering the mirror system as not a passive simulation mechanism, but as supporting predictive mechanisms, may help further research on action observation. Our experiment used music as a tool to explore this question. It is possible that music is particularly special in its multimodal, sequential nature. It remains to be tested whether different kinds of sensory prediction also involve active prediction in the motor system, or if other contexts (i.e., novel actions) may invoke a more passive, resonant role. Future work is warranted to investigate other kinds of sensory prediction and involvement of the motor system in these domains.

## Author Contributions

CG, MI and RB initiated this work following workshops for the UC MERCI (Music experience research community initiative) group. All of these authors contributed substantially to this work, and approved it for publication. CG performed the experiments, completed the analyses and made the graphs. RB and MI contributed both to the experiment design and to the intellectual components of the work.

